# Isolation to Extinction: Landscape Fragmentation and Connectivity in Sloth Bears of the Indian Terai

**DOI:** 10.1101/2025.09.27.678944

**Authors:** Divyashree Rana, Judi Clarissa Samjith, Uma Ramakrishnan

**Affiliations:** National Centre for Biological Sciences, Tata Institute of Fundamental Research, Bangalore, India; Department of Biotechnology, Manipal Institute of Technology (MIT), Manipal Academy of Higher Education (MAHE), Manipal, 576104, Karnataka, India

**Keywords:** Sloth bear, fragmentation, inbreeding, isolation, local extinction, non-invasive samples, non-model organisms, peripheral populations, SNPs from feces

## Abstract

Habitat fragmentation is a major global threat to biodiversity, leading to small and isolated populations. Fragmented landscapes can reduce connectivity, erode genetic diversity, and increase inbreeding risk, potentially contributing to local extinction. Using a landscape genetics approach, we investigated sloth bear (*Melursus ursinus*) populations in India’s Central Terai landscape - a human-dominated, multi-use region that exemplifies the challenges of conserving wildlife populations in highly fragmented landscapes. Using non-invasively collected faecal samples (n=209), we identified 66 individuals from 964 genome-wide SNPs generated via ddRAD sequencing. Spatial genetic structuring was not strictly congruent to the protected areas and genetic clusters showed low genetic diversity (mean H_E_ = 0.156 ; mean H_O_ = 0.079) with a heterozygote deficit (mean F_IS_ = 0.323). Further, landscape genetic analyses revealed agriculture as the primary barrier to gene flow across the landscape, followed by proximity to water. Our findings establish the first genetic baseline for sloth bears in this region and suggest that historical population structure and contemporary fragmentation may jointly influence genetic connectivity and diversity across the landscape. We demonstrate the unique utility of non-invasive samples and genome-wide SNP based analyses for poorly studied species of conservation concern, despite experimental and bioinformatic challenges. Our novel insights underscores the importance of such genetic monitoring in fragmented landscapes.

## 1. Introduction

Fragmentation of habitats not only reduces the overall availability of suitable area but also hinders animal movement, thereby impeding their access to mates and resources (Francis et al., 2016). Predominantly driven by anthropogenic factors such as urbanisation, agricultural expansion, and industrialisation, it is recognised as a major threat to biodiversity globally (Fahrig, 2003). However, responses to human-modified landscapes are species-specific; while some adaptable species exploit these novel environments, for many they create isolated patches within unsuitable matrices that limit dispersal (LaPoint et al., 2015). These small and isolated populations may exhibit increased inbreeding (Frankham, 1998), reduced allelic richness (Allendorf et al., 2024) and heightened demographic stochasticity (Araya-Ajoy et al., 2025), ultimately lowering fitness and evolutionary potential (Allendorf, 1986). These effects are particularly severe for species with limited dispersal abilities and large home ranges (Crooks, 2002). Conservation scenarios such as the inbreeding depression of the Florida panther prior to genetic rescue (Johnson et al., 2010), the prolonged isolation and collapse of Isle Royale’s gray wolves (Hedrick et al., 2014) and the unusually high frequency of a rare recessive trait in the melanistic tigers of Similipal Tiger Reserve (Sagar et al., 2021) provide documented evidence of the genetic and demographic consequences faced by small, isolated populations. Collectively, these cases underscore the critical importance of maintaining habitat connectivity as a conservation priority to ensure long-term population viability.

Given the conservation concerns associated with small and isolated populations, it is essential to understand the role of both natural and anthropogenic fragmentation in maintaining functionally connected populations. Assessing how landscape features influence genetic connectivity allows us to identify habitat features that influence species’ movement to design connectivity corridors (Anderson et al., 2015; Modi et al., 2025). Identifying and prioritizing such areas for conservation action is critical for effective management and long-term population viability. Recent advances in analytical approaches to optimize resistance surfaces and predict functional connectivity have improved our understanding of species-specific landscape permeability (Peterman, 2018; Shirk et al., 2018). These tools have helped evaluate the effectiveness of the existing protected areas and corridors (Naidoo et al., 2018) as well as prediction of how future land use changes may affect dispersal and movement (van Strien et al., 2014). When integrated with empirical genetic data, such models can uncover cryptic isolation patterns (Wasserman et al., 2012) and detect functional corridors (Sharma et al., 2013), offering particular value for wide-ranging and elusive taxa.

Parallely, advancements in DNA extraction and sequencing technologies have expanded the potential of non-invasively collected faecal, hair or saliva samples to infer population-level genetic parameters, including structure, connectivity and diversity, without directly handling individuals (Taberlet et al., 1999; Andrews et al., 2018). These DNA sources have been particularly valuable for elusive and endangered species but require careful optimization to address the low quantity and degraded quality of DNA typically obtained (Bourgeois et al., 2019). Traditionally, capillary-based fragment analysis of hypervariable microsatellites has been employed for such samples; however, these markers are limited by their number, prone to genotyping errors, and often suffer from low reproducibility (Estoup et al., 2002; Sittenthaler et al., 2020). Amplicon-based sequencing using standardized marker panels that target short DNA fragments has emerged as a reproducible and more robust yet species-specific alternative (De Barba et al., 2017; Natesh et al., 2019; Rana et al., 2024). Beyond targeted panels, reduced-representation genomic approaches such as Restriction site-Associated DNA sequencing (RAD-seq) enable simultaneous discovery and genotyping of hundreds of SNPs, but their application from non-invasive samples remains limited (but see Tyagi et al., 2022; Balmori de la Puente et al., 2023; Scott & Kovach, 2024). With the increasing accessibility of these genomic approaches, non-invasive sampling when processed rigorously offers a powerful means to overcome the logistical challenges of direct monitoring and to generate robust insights into the genetic processes shaping populations in fragmented landscapes.

The Central Terai Landscape (CTL) in India, an altered multiuse landscape characterized by high human density, anthropogenic pressures, and rich biodiversity, exemplifies the challenges of conserving a wide-ranging species in shared spaces (Thapa et al., 2021). The sloth bear *(Melursus ursinus),* a large carnivore within this landscape, is now largely restricted to protected areas. With extensive agricultural expansion in the landscape since the 1970s, today the remnant habitats are embedded within a matrix of agriculture and settlements. Hence, understanding the functional connectivity of these sloth bear populations is critical because they represent an important component of the transboundary population across the fragile floodplain shared by India and Nepal. To address this, we used a ddRAD-based approach applied to non-invasively collected faecal samples to investigate population structure, connectivity and its drivers in the landscape. Specifically, we aimed to (i) optimize a non-invasive genetic protocol for monitoring sloth bears, (ii) investigate if the sloth bear populations are genetically structured across the landscape, (iii) assess whether protected areas mirror the genetic clustering or harbour genetically similar individuals, and finally (iv) map functional connectivity corridors between these protected areas. Given the combined pressures of habitat fragmentation and range-edge effects, we predicted that sloth bear populations in the CTL will exhibit reduced genetic diversity and elevated genetic differentiation across protected areas, reflecting the impact of intense landscape modification.

## 2. Methods

### 2.1. Study area, focal species, and sampling

Non-invasive sampling was conducted across the core and buffer areas of Pilibhit Tiger Reserve (PBT), Dudhwa National Park (DDW), Kishanpur Wildlife Sanctuary (KIS), and Katarniaghat Wildlife Sanctuary (KAT) in the CTL of India, during the dry winter months (December - March) of 2022 and 2024. These protected areas range in size from approximately 200-800 km² and are separated by distances of roughly 15-20 km from adjacent parks. The broader study landscape spans over 8,000 km² and comprises a mosaic of natural species habitat largely restricted to protected areas (<2,500 km² in total) embedded within a matrix of human-modified landscapes. This region, part of the larger Terai Arc Landscape (TAL) spanning from southern Nepal to the northern regions of India, is a biologically and ecologically rich landscape for sloth bears amongst a wide range of flora and fauna (Ram et al., 2021). However, extensive habitat fragmentation and forest loss driven by human population expansion, agriculture, and infrastructure development have increased human-wildlife conflict and contributed to population declines of several species (Thapa et al., 2021), making this landscape well suited for investigating the consequences of habitat fragmentation.

This region forms the northernmost extent of the range of the sloth bear (*Melursus ursinus*), a large omnivore endemic to the Indian subcontinent with India harbouring 90% of its global population. Listed as vulnerable on the IUCN redlist, their populations are isolated across the country, primarily due to habitat fragmentation (Dharaiya et al., 2017). There is a lack of studies revealing key ecological insights about the species including home ranges and movement, specially in modified landscapes. In the adjoining Terai populations in Nepal, the average home ranges are around 11 km², larger for males (14.4 km²) than females (9.4 km²). Overall, sloth bear home ranges are relatively small and flexible, reflecting their dependence on patchily distributed food resources and habitat types (Ratnayeke et al., 2007). Although we could not find records of estimated dispersal distance, it is calculated to be around ∼13 km based on the recent global mammal trait database (Soria et al., 2021). In many parts of the country, sloth bears face conflict with humans and are known to move in human-modified urban and agricultural landscapes (Singh et al., 2018; Desai et al., 2025). However, unlike other populations in India, sloth bears in this transboundary landscape with Nepal are restricted to protected areas with lack of reports of movement in human modified areas (Sadadev et al., 2024). The limited distribution of the species and presumed lack of movement across protected areas, despite being within the estimated dispersal limits of the species, raise concern and underscore the need to investigate the genetic population structure and barriers to genetic connectivity. Hence, suspected sloth bear faecal samples were collected non-invasively using sterile swabs and preserved in Longmire’s buffer (Longmire et al., 1997) to understand population connectivity across the landscape.

### 2.2. Generating genetic data and identifying species

DNA was extracted using Qiagen Blood and Tissue Extraction kit following the manufacturers protocol and concentrations were measured using Qubit Fluorometer 4.0 (*Invitrogen*). A two-step approach was adopted to genetically confirm the species origin of the suspected faecal samples to account for poor-quality of collected samples and lower amplification rate of species-specific markers. The extracted DNA was first screened with a species-specific *COXII* marker (Thatte et al., 2018) and extracts that failed to amplify were subsequently tested with a more generic *16S rRNA* marker (Mukherjee et al., 2018) (protocol details in Supplementary Material S1). The *16S* amplified products were sequenced in the in-house Sanger sequencing facility and sequences were matched to the NCBI genetic repository (>97%) for robust species identification. Extracts that failed to amplify using the *16S rRNA* primers were concluded to be poor-quality and discarded from downstream analyses.

Owing to the presence of prey and bacterial contamination, faecal samples have low host DNA concentration. Thus, the samples genetically identified as belonging to sloth bears were enriched using methyl-CpG-binding domain (MBD) proteins to maximise host DNA concentration (Chiou & Bergey, 2018). The methylated bead-based enrichment method works on the biological principle that eukaryotic DNA is more heavily methylated compared to microbial DNA, hence the magnetic beads coated with MBD2-Fc protein preferentially binds to methylation-rich vertebrate DNA (Wood & Zhou, 2016). For DNA enrichment, NEBNext Microbiome DNA enrichment kits were used (Feehery et al., 2013), for which MBD2-Fc magnetic beads were prepared by the conjugation of MBD2Fc protein and protein A magnetic beads (details in Supplementary Material S2). Prior to enrichment, the extract volume was reduced from 200 to 40uL using the CentriVap DNA concentrator (*Labconco*) to concentrate the DNA.

Post enrichment, the extracts were processed using double digested library preparation protocol (Peterson et al., 2012), while incorporating some modifications specific to low host DNA concentration samples (similar to Tyagi et al., 2022). A combination of *Sph1* and *MluC1* restriction enzymes was used for digestion at 37°C for 3 hours followed by a 20 minutes heat-kill at 65°C, followed by adapter ligation using T4 DNA Ligase at 23°C for 2 hours and 10 minutes at 65°C. Ligated products were purified to remove <100 bp fragments, followed by a 16-cycle indexing PCR (98°C for 30s, 65°C for 30s, 72°C for 30s and 72°C for 5min). Post indexing, fragments ranging from 200-500 bp were retained during purification (0.5X and 1X) with in-house prepared carboxyl group-coated SPRI beads. Individual sample libraries were quantified using a Qubit Fluorometer and their fragment size distribution was evaluated using the Agilent Tapestation (detailed library preparation protocol in Supplementary Material S2).

DNA concentrations were measured after each processing step - i. extraction, ii. enrichment, and iii. library preparation - and products with unquantifiable DNA concentration were discarded. The remaining sample libraries were pooled in an equimolar fashion to generate comparable reads across varying sample concentrations. The pooled library was ultimately purified to retain fragments ranging from 200 to 500 bp (Figure S2.1), and was sequenced on the NovaSeq platform.

### 2.3. Calling and filtering variants

Demultiplexed reads were quality checked using *FastQC* and compiled to a single *.html* report using *MultiQC v1.26* (Ewels et al., 2016). As the reads were generated from faecal samples, contamination with prey as well as bacterial DNA was a major concern despite the enrichment. Hence, the raw reads were trimmed for adapter sequences and low quality bases, using *Trimmomatic v0.36* (Bolger et al., 2014), and then mapped to the reference genome of the closest related species - sun bear *Helarctos malayanus* (GCA_028533245.1) (divergence time of ∼3.5 million years with the last common ancestor) (Kumar et al., 2017), in absence of a sloth bear reference assembly at the time of analyses (but see Wooldridge et al., 2025), using *BWA-MEM v0.7.17* (Li, 2013). The bam files, retaining only the mapped reads, were then converted to a .fastq file using *BEDTools v2.30.0* (Quinlan et al., 2010) to be used for downstream processing.

The filtered reads were then processed using the *ipyrad* bioinformatic pipeline to conduct de-novo local assembly and analyze the ddRAD sequence data (Eaton & Overcast, 2020) using default parameters to generate a Variant Call Format (VCF) file with SNPs. The filters ensured a minimum depth of six bases, 85% clustering threshold, and retention of loci present in a minimum of four samples. Additionally, reads with more than five low quality bases, phred score of less than 33, or sequence length less than 35 were removed (more details in Table S3.1). In an attempt to improve subsequent coverage and geographical representation of samples, the ones that yielded little sequencing data (> 10 MB) and were filtered out during variant filtering were sequenced once more and raw data from the same samples was merged, before repeating the filtering process.

Reduced-representation sequencing approaches such as ddRAD may experience allele or locus dropout due to restriction-site polymorphisms or uneven sequencing coverage (Gautier et al., 2013). To minimize such effects, the output VCF file containing the variants was further filtered using *vcftools v0.1.17* (Danecek et al., 2011), an approach utilised by other similar studies (Andrews et al., 2016; Tyagi et al., 2024; White et al., 2026). Genotypes with sequencing depth below 6x or above 1000x were treated as missing, and variants were subsequently filtered to retain biallelic sites with a minor allele count (MAC) >3, thereby excluding variants supported by very few allele copies.. Lastly, samples with over 60% and loci with over 40% missing data were dropped to remove poor quality data and reduce overall missingness. The data generation, filtering and its subsequent population genetic analyses has been summarised visually in Figure 1. Although our analyses is based on reference genome of a closely related species — sunbear, we repeated the above described variant calling and filtering along with measurement of comparable genetic diversity estimates using the later published sloth bear reference genome (BioProject PRJNA1284380) as well to validate our approach (see Table S4.1).

**Figure 1:**
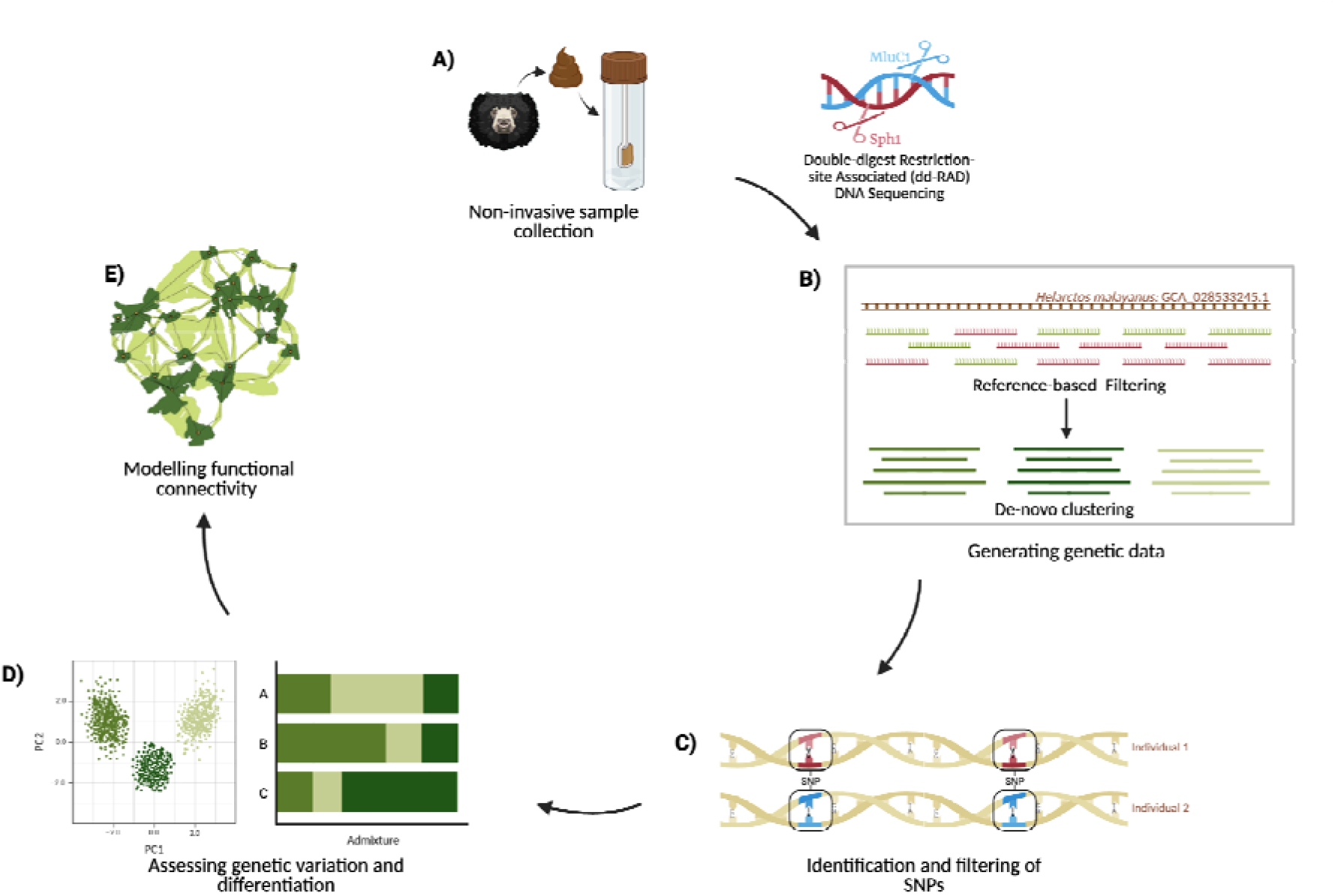
Schematic explaining the methodological workflow. A) Non-invasive faecal sample collection of sloth bears using sterile swabs; B) Genetic data generation workflow (Preparation of ddRAD-sequencing libraries, mapping of sequenced reads to a closely related reference genome to filter sequences and de-novo clustering of filtered sequences); C) Identification and filtering of SNPs across individuals; D) Assessment of genetic variation and differentiation across sampled protected areas; E) Identification of gene flow barriers and modelling functional connectivity across the landscape *Created in BioRender*.

### 2.4. Assessing genetic variation and differentiation

Potential recaptures or duplicate samples in the dataset could result in false inferences of genetic clusters, hence the genetic relatedness between samples was assessed to identify unique individuals and remove any individual recaptures using *KING: Relationship Inference Software v2.3.1* (Manichaikul et al., 2010). To investigate the extent of genetic mixing among individuals and identify shared ancestries, genetic structure was assessed using *ADMIXTURE v1.3.0* (Alexander et al., 2015) without any *a priori* groupings. Using .bed files generated in *PLINK v1.9* (Purcell et al., 2007), the algorithm was run for a range of clusters (K = 1-10), with 25 independent replicates performed for each K. The optimal number of clusters was determined using cross-validation error. Replicate runs were summarized using *CLUMPAK* (Kopelman et al., 2015), which aligns cluster assignments across runs using the algorithm implemented in CLUMPP and generates consensus ancestry coefficients for visualization. Individuals were assigned to the genetic cluster for which they had the highest ancestry membership coefficient, and these assignments were used to define genetic clusters for subsequent estimates of genetic diversity. The level of genetic variation within identified genetic clusters was then assessed by calculating the observed (H_O_), Nei’s unbiased expected (H_E_) heterozygosity, and population-level inbreeding coefficient (F_IS_) using the “*basic.stats*” function of the *hierfstat v0.5-11* package (Goudet, 2005) in R. Lastly, sensitivity of individual-level inbreeding coefficients (F) to genotype missingness and underlying population structure was assessed, where individual F was estimated using the likelihood-based “*inbreeding*” function in the *adegenet v2.1* package (Jombart, 2008) in R. This allowed us to assess the extent to which individual-level F estimates may be influenced by incomplete genotyping and the choice of underlying population structure, a challenge inherent with degraded non-invasive samples in fragmented landscapes.

Additional to *a priori* genetic population structure, given the small spatial scale of the study and local management reliant conservation planning for the species, our next objective was to understand genetic differences between samples collected from different management units — the protected areas. To assess whether genetic clusters corresponded to geographic management units, individuals were assigned to sampled protected areas as well as the resulting genetic clusters. To explicitly visualize genetic differentiation among geographically clustered individuals, a supervised classification step was implemented in Discriminant Analysis of Principal Components (DAPC) using the *adgenet* package in R. Individuals were assigned to groups based on predefined geographic labels corresponding to the protected areas from which they were sampled. The optimal number of principal components retained for DAPC was determined using the cross-validation procedure implemented in the “*xvalDapc*” function in *adegenet*. Based on assignment success across 100 replicates, 15 principal components were retained for the final analysis. Lastly, to compare the degree of genetic differentiation between protected areas, the fixation index (F_ST_) was computed using the “*pairwise.WCfst()*” function of the *hierfstat* package, allowing inference about the degree of connectivity between them.

### 2.5. Modelling functional connectivity across landscape

Pairwise genetic and geographic distances were correlated to understand the effect of Isolation By Distance (IBD) in shaping the spatial population structure using mantel test from the *ade4* package in R (Dray et al., 2007). Here, genetic distances were computed as the proportion of allelic dissimilarity (Prevosti’s absolute genetic distance) using the “*diss.dist”* function in the *poppr v2.9* package in R (Kamvar et al., 2014), setting percent = TRUE to standardize for the number of loci. The pairwise geographic distances were calculated using the “*spDists”* function in the *sp v2.1* package in R (Pebesma & Bivand, 2005).

To further understand the influence of landscape features on observed spatial population structure for the species, a non-linear landscape genetic approach was implemented. This involves parametrization of resistance surface for each landscape feature determining the resistance offered by the latter to movement of the species using pairwise genetic distances. Based on sloth bear ecology, landscape properties, and published literature, four uncorrelated landscape variables (distance to water, agriculture land cover density, enhanced vegetation index, and distance to human settlement) relevant to understanding the connectivity of sloth bear populations were selected (Table 1 and Figure S5.1) (Dutta et al., 2015; Thatte et al., 2020; Malik et al., 2023). All variables were reclassified and recalculated as explained in Table 1 and mapped in Figure 4B. Distance layers were generated using the *“Proximity (Raster Distance)”* function in QGIS with a binary target layer as the input raster. As studies have shown that species-landscape interactions are scale-dependent (Cushman & Landguth, 2010; Suárez-Castro et al., 2018), all variables were processed at four spatial scales (1 km, 2 km, 5 km, and 10 km) with a moving window focal mean using the *terra v1.7* package in R (Hijmans et al., 2022). Notably, this moving window analysis allows retaining the finest “resolution” (or pixel size) for each raster (1 km) while changing the “scale” of analyses (see Figure S5.3), theoretically mimicking the scale at which the species are interacting with the environment (Peterman et al., 2018). Hence, this processing of the rasters at different scales, allows making multiscale composite rasters at the end based on optimized scale-selected rasters (further details can be found in Supplementary Material S5).

**Table 1.** Description and details of landscape variables used in the study.

| Variable | Description | Source | Details |
| --- | --- | --- | --- |
| Distance to water | Distance from the centre of each pixel to the center of the nearest permanent inland water | Google Earth Engine<br>(ee.ImageCollection("GLCF/GLS_WATER")) | Binary water layer based on category 2: "water" |
| Agriculture landcover density | Percentage of area occupied by agriculture landuse | Google Earth Engine<br>(ee.ImageCollection("ESA/WorldCover/v100")) | Per pixel density for category 40: "agriculture" |
| Enhanced vegetation index (EVI) | Satellite-derived metric for vegetation density correcting for atmospheric noise | Google Earth Engine<br>(ee.ImageCollection("MODIS/MOD09GA_006_EVI")) | Raster scaled by a factor 0.0001 |
| Distance to human settlement | Distance from the centre of each pixel to the center of the nearest to human settlement | Google Earth Engine<br>(ee.ImageCollection("JRC/GHSL/P2023A/GHS_BUILT_C")) | Binary settlement layer generated by combining categories 12-25 (builtup area) |

A multi-scale multi-surface optimization approach was used to understand the non-linear effect of landscape features in governing gene flow, and to create a composite resistance surface using *resistanceGA v4.0* package in R (Peterman et al., 2018). This was based on a mixed effect linear mixed effects model fitted to the genetic data with a maximum likelihood population effects parameterization (MLPE). The MLPE framework effectively accounts for the non-independence and shared covariance structure inherent to pairwise distance matrices, and it has been demonstrated as an optimal approach for landscape genetic inference (Shirk et al., 2018; Peterman et al., 2019).

As sloth bears are primarily solitary in nature, individual-based analysis without any *a priori* groupings was implemented to understand landscape effect on gene flow across sampled individuals. Genetic distances were calculated between unique individuals using the “*diss.dist*” function as explained above. The rasters for the landscape variables were converted to *ascii* format using the *terra* package in R. With genetic data as the response and effective cost distance within each landscape variable as predictors, single surface models were optimized using pairwise least-cost paths via the *gdistance v1.5* package (Van Etten, 2017), and executed with default parameters to attain lowest Akaike Information Criterion (AIC) values within a maximum of 1000 iterations. Hence, at every scale, each landscape variable was optimized to identify the transformation type, shape of the transformation, and the maximum resistance offered using the “*SS_optim”* function in the R*esistanceGA* package in R. Therefore, using univariate optimization, scale and best-suited model parameters for each landscape variable were identified. These optimized single surface parameters (Figure S5.3) were used to understand the non-linear influence of landscape variables on genetic population structure.

These optimized parameters for each scale-selected raster variable (Figure S5.3) were further used to generate composite surfaces with all combinations of the four variables to optimize the composite landscape resistance layer (Figure S5.4) using the “*MS_optim”* function in the *ResistanceGA* package in R. The optimized composite multisurface raster layer with least AIC value was used to model functional connectivity across the landscape. Using the optimized multisurface resistance layer and protected area node raster, the connectivity was modelled for sloth bears using the pairwise modelling mode with default parameters in *Circuitscape 4.0* software (McRae et al., 2009). Finally, top 5 and 10% quantile values were used to identify potential critical corridors between the surveyed protected areas (Poor et al., 2024; Fonda et al., 2026).

## 3. Results

### 3.1. Genetic data generation and species identification

Out of the 209 suspected sloth bear samples collected, 189 samples were successfully identified using the two-step species-specific (n=126) and *16s* (n=63) primer amplification steps. All confirmed sloth samples were subsequently enriched, out of which 111 samples with quantifiable post-enrichment concentrations were prepped and added to the final library pool owing to variable DNA quality and concentration. Post sequencing, poor-quality samples with little or no sequencing reads (<10 MB) were dropped. A raw VCF file identifying a total of 697,579 SNPs across 94 individuals was generated using the *ipyrad* pipeline, however it had very high missingness. Hence, post filtering and merging resequenced samples with few reads, a final filtered vcf containing 964 SNPs across 66 samples was retained (see Table 2, Figure S1.1).

**Table 2:** Progression of sloth bear samples through species identification, enrichment, library preparation and SNP recovery.

| Protected Areas | Sampling effort (in km) | Collected Samples | Confirmed Sloth Bear Samples | Enriched samples | Unique Individuals |
| --- | --- | --- | --- | --- | --- |
| DDW | 975 | 68 | 58 | 26 | 10 |
| PBT | 1064 | 85 | 77 | 57 | 41 |
| KIS | 510 | 56 | 54 | 28 | 15 |
| KAT | 571 | 0 | 0 | 0 | 0 |
| **Total** | **3120** | **209** | **189** | **111** | **66** |

### 3.2. Genetic variation and differentiation

All samples successfully genotyped represented unique individuals with no recaptures (Figure S4.1). The ADMIXTURE plot illustrates admixed individuals with genetic structuring based on individual ancestry composition (Figure 2B). The optimal K based on least cross-validation (CV) error across ADMIXTURE runs was identified as four (Figure 2A). At K=3, DDW individuals show predominantly a single ancestry. At the optimal K=4, individuals from DDW exhibited dominant ancestry from a single cluster (dark green) with two probable admixed individuals. KIS individuals also show a similar pattern with a dominant ancestry of a single cluster (pink) with few admixed individuals sharing the genetic cluster (light green) dominant in PBT. Conversely, individuals from PBT display highly mixed ancestry with proportions from all four clusters (Figure 2B). The spatial genetic structure map, with pie charts representing admixture proportions of individuals across the landscape, further emphasises the spatially local dominance of certain clusters (Figure 3). Based on relative highest membership, each individual was segregated into a respective genetic cluster or population within the identified four optimal clusters. Similar to the visual representation in Figure 3, genetic populations did not mirror geographic management units with unequal genetic distribution across sampled protected areas. DDW individuals primarily belonged to the genetic cluster K1 (n = 9) while the KIS individuals belonged to K4 (n = 11). However, PBT individuals belonged to all four genetic clusters, but most prominently K1 (n = 18) and K3 (n = 14), with K2 as the smallest overall genetic cluster (Table S4.2). Within identified genetic populations, a low mean observed heterozygosity (H_O_) of 0.079 (range: 0.041 - 0.136) and a mean expected heterozygosity (H_E_) of 0.156 (range: 0.079 - 0.297) was observed. Notably, a positive inbreeding coefficient (F_IS_) was observed for all identified genetic clusters with a mean of 0.323 (range: 0.170 - 0.442) (Table S4.3). Based on the sensitivity analyses, positive individual heterozygosity deficits were not associated with genotype missingness (Figure S4.3), and were retained even after accounting for underlying genetic structure (Figure S4.4).

**Figure 2:**
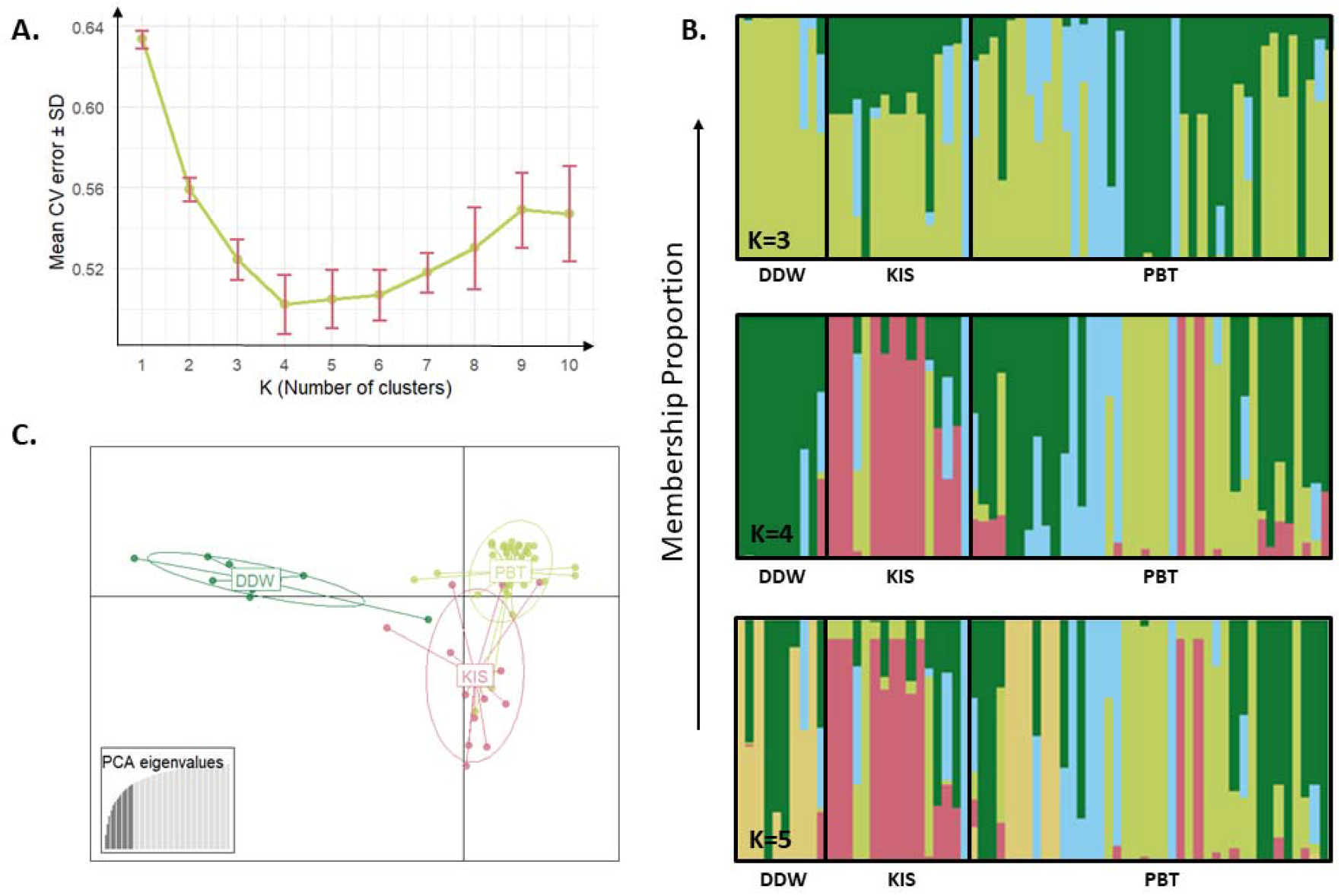
Investigating population structure of sloth bear populations in CTL. A. Identifying optimal K based on cv error across admixture runs B. Admixture plot showing membership proportions for individuals under different optimal K across protected areas C. Discriminant analysis of principal components representing variation across groups based on supervised clustering across protected areas.

**Figure 3:**
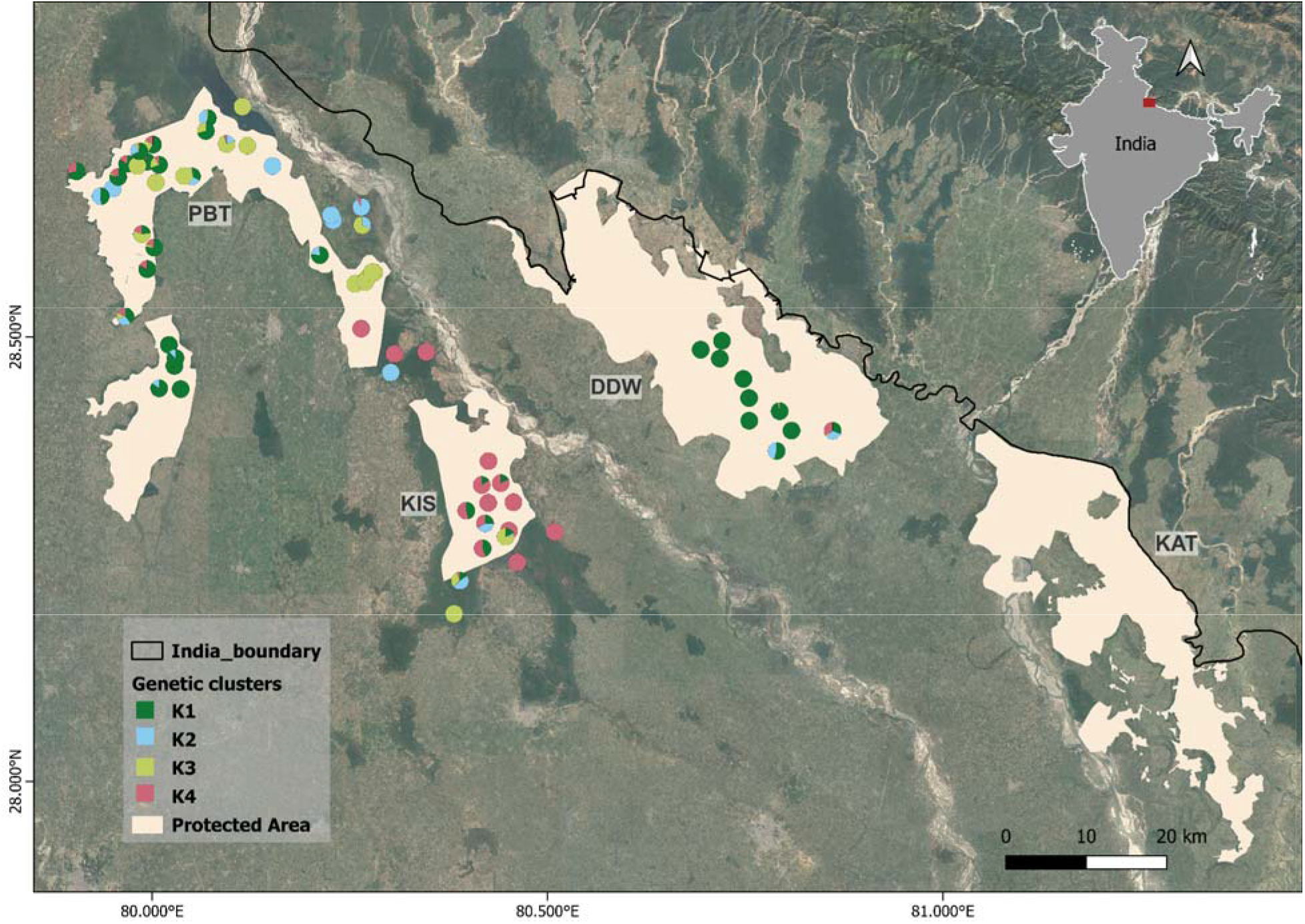
Geographical distribution of genetic clusters of sloth bear populations based on the identified optimal K=4 across the sampled protected areas. The red box on the India map (top right corner) marks the location of the study area.

**Figure 4:**
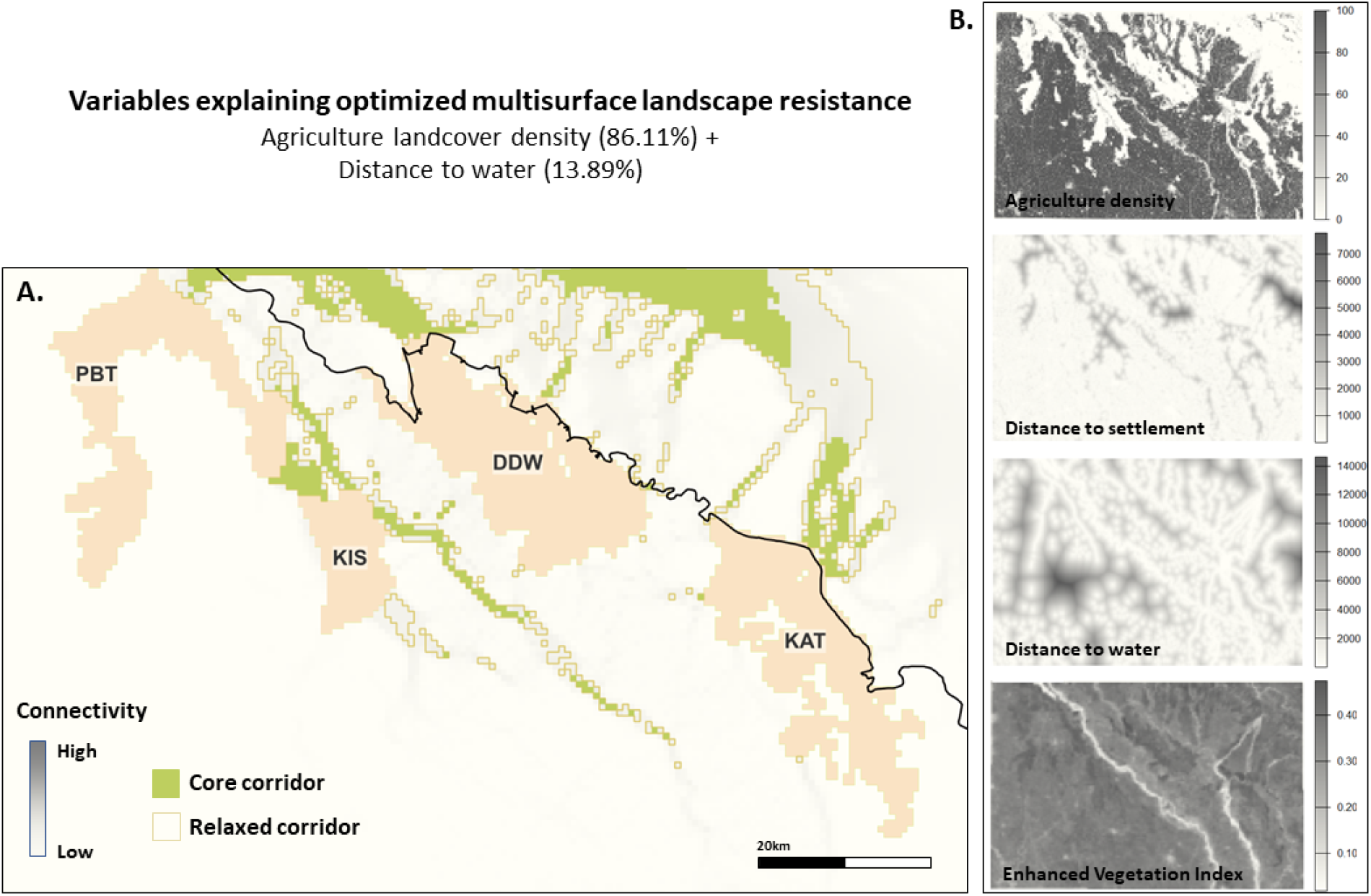
Predicted connectivity map based on multisurface optimization of landscape variables. A. Circuit theory based connectivity map based on optimized composite resistance surface. B. Input landscape variables (n=4) with 1 km resolution used to build an optimized composite resistance surface.

In line with the AMDIXTURE findings, the protected area labelled supervised clustering in the DAPC plot (Figure 2C) reveals a distinct separation between individuals of DDW which formed a non-overlapping cluster, compared to those of PBT and KIS in which partial overlap can be seen. Pairwise F_ST_ values across protected areas show a similar trend with highest differentiation between DDW and KIS (F_ST_ = 0.30), followed by DDW and PBT (F_ST_ = 0.14), and least differentiation between PBT and KIS (F_ST_ = 0.09) populations.

### 3.3. Population connectivity patterns

A weak positive yet statistically significant correlation (r = 0.09; p = 0.042) was observed between genetic and geographic distances using the Mantel test. Visual inspection of the data density distribution reveals a continuous, reasonably linear trend without major non-linear thresholds or distinct localized clusters (see Figure S4.5). The density is heavily concentrated at intermediate to long spatial scales (∼50-85 km) with a broad vertical distribution of genetic distances (0.4 to 0.8) at any given geographic interval.

The univariate optimization revealed variable optimal scales of effect for different landscape variables, smaller for agriculture density (1 km) and distance to water (2 km) while larger optimal scales for distance to settlement (5 km) and EVI (10 km) (Table S5.1). All landscape variables showed different relationships with genetic distance after single surface optimization. Agriculture showed a positive exponential relation, with resistance to movement increasing rapidly in areas with over 80% agriculture landcover density. With increasing distance to water, the resistance to gene flow decreased sharply with almost no resistance after 6 km. On the other hand, resistance decreased with increasing values of EVI. The EVI in the landscape had a narrow range from 0.2 to 0.4, and the optimized parameters showed low resistance to movement in areas with denser vegetation. Lastly, distance to settlement showed a positive exponential relation with high resistance values after 1 km (Figure S5.2).

The optimized composite multisurface resistance layer had two landscape variables influencing genetic structure (Table S5.2). Agriculture land cover density was the most important and dominant variable explaining the population structure with 86.11% variable contribution, followed by distance to water with 13.89% contribution. The modelled connectivity map showed overall weak connectivity across the landscape (Figure 4A). The identified critical connectivity hotspots or corridors were primarily between PBT and KIS, showing maximum connectivity between them whereas there were fragmented corridors between DDW and KIS and none between DDW and KAT. Transboundary connectivity with Nepal was also highlighted.

## 4. Discussion

This study demonstrates how integrating robust genetic data from non-invasive samples with complex landscape genetic approaches can yield critical insights into the genetic variation and movement ecology of wide-ranging, elusive species inhabiting fragmented landscapes. Focusing on sloth bears in the Central Terai Landscape (CTL), a multi-use and populated landscape of India, our analyses reveal how recent fragmentation accentuating probable historical processes shapes genetic erosion to result in local extinction. We detected geographically structured populations with evidence of admixture across protected areas. While the genetic populations did not correspond completely with geographical boundaries limited to protected areas, identified genetic populations exhibited low genetic diversity measures with heterozygote deficit. Further, non-linear landscape genetic analyses identified agricultural density as the primary barrier to gene flow across the landscape. Together, these findings suggest that sloth bears in this landscape possibly have small populations and face higher risk of inbreeding making them genetically vulnerable. Our study presents a timely example of population isolation driving local extinction of a large carnivore, while providing process driven explanation for the event. With increasing habitat fragmentation globally, such spatial genetic insights are critical for endangered species, especially for peripheral populations that are often characterised by historical bottlenecks and founder effects.

### 4.1. Non-invasive genomic approaches enable multiscale insights

Our study represents one of the few applications of non-invasive genomic methods using hundreds of SNPs to investigate population genetics of a large carnivore in India, and the first genome wide assessment of sloth bears. Working with faecal samples posed inherent challenges of degraded DNA, but optimized protocols significantly improved genotyping success. Species identification using a sloth bear-specific primer yielded only 60.3% success rate (n = 126), lower than Thatte et al. (2018), who reported 87%. However, our modified two-step approach, aimed to increase amplification success and downstream sample retention, achieved 90.4% success (n = 189), demonstrating the power of targeted protocol optimization.

A major limitation for many non-model species, including sloth bears, is the lack of a readily available species-specific reference genome (Brandies et al., 2019). This gap, particularly pronounced in biodiverse yet under-resourced regions of the Global South (de Jong et al., 2024; Pitogo, 2025), constrains conservation genomic applications. Multiple international initiatives such as the Earth BioGenome Project (Lewin et al., 2018) and the European Reference Genome Atlas (McCartney et al., 2024) aim to address this to aid conservation monitoring. In our study, mapping non-invasive reads was complicated with a lack of reference genome, especially from wild Indian populations, as well as contamination of faecal samples with prey and microbes (Formenti et al., 2022). We addressed these challenges through a combined strategy of reference-based filtering using the genome of closely related species, followed by *de novo* clustering, retaining ∼35% of samples for robust downstream analysis. This approach enabled reliable individual identification and assessment of fine-scale genetic structure and diversity within populations, demonstrating the feasibility of robust non-invasive genetic monitoring with hundreds of genome-wide SNPs even in species lacking reference genomes. Additionally, with the later availability of a sloth bear genome, we found similar genetic diversity estimates as well as sample retention post filtering obtained using both species-specific as well as closely related species reference genome, hence lending further support for our approach. Although the choice of using sun bear reference genome didn’t influence our findings pertaining to spatial population structure at the relevant scale, we strongly believe that increasing availability of high quality reference genomes would permit for adaptive as well as historical understanding of populations (Brandies et al., 2019), which was beyond the scope of our study.

### 4.2. Population genetic patterns suggest historic bottlenecks

We detected spatially structured populations with four partially admixed genetic clusters spread across protected areas (Figure 2B). DDW formed the most homogeneous group, while KIS and PBT exhibited increasing proportions of genetic admixing. These patterns are consistent with geography - gene flow appears more likely between the structurally contiguous PBT-KIS protected areas with continuous natural habitat, than between either of them and DDW, which is more isolated, specifically by presence of major river systems in between. Pairwise F_ST_ estimates across sampled protected areas also supported this interpretation, with lower differentiation between PBT-KIS relative to DDW.

Despite varying degrees of admixture across sampled areas, overall unbiased observed (0.079) and expected (0.156) heterozygosity were low with consistent heterozygote deficit, indicating loss of genetic variation. Multiple plausible drivers could be contributing to these findings. Our sensitivity analyses revealed that positive individual-level F estimates cannot be explained by genotype missingness or Walhund effect in structured populations alone, however the influence of allelic dropout cannot be ruled out entirely. Nevertheless, these genetic diversity estimates could also be suggestive of small population size and historical bottlenecks, often associated with populations at range edge. The central marginal hypothesis predicts reduced genetic diversity and heightened differentiation toward species’ range edges (Eckert et al., 2008), and these sloth bear populations present an example of such peripheral populations. Microsatellite-based studies of sloth bear populations in central and western India report higher heterozygosity estimates (Dutta et al., 2015; Gomes et al., 2024; Sharma et al., 2013; Thatte et al., 2020). Although absolute heterozygosity values are not directly comparable across marker types (Pérez-González et al., 2023), these studies report observed heterozygosity comparable to or exceeding expected heterozygosity estimates, indicating little or no heterozygote deficit. In contrast, our SNP dataset shows observed heterozygosity relatively lower than expected heterozygosity estimates, indicating a positive heterozygote deficit, similar to the findings from a recent study on adjacent Nepal populations (Paudel, 2026). Despite the risk of allelic dropout inflating these estimates, we believe our extensive filtering criteria following other similar studies (Andrews et al., 2016; Tyagi et al., 2024; White et al., 2026), and the inferences from the sensitivity analyses presents evidence that inbreeding risk does exist in these likely small populations. Knowing population sizes could help bolster some of these claims but unfortunately for unmarked species like sloth bears, population sizes are virtually unknown across its range. Notably, despite extensive and comparable sampling effort in DDW and PBT (Table 2), we identified much fewer individuals in DDW which could be suggestive of its smaller population size, further contributing to witnessing lower genetic diversity and higher differentiation measures. This pattern suggests overall reduced genetic diversity and potential demographic stress in the study population.

Populations that have undergone historical bottlenecks or experienced long-term fluctuations in effective population size often retain the genetic signatures of drift, including reduced heterozygosity, elevated inbreeding, increased genetic load, and loss of rare alleles (Abascal et al., 2016; Battilani et al., 2025). Such erosion of genetic variation constrains adaptive potential and amplifies extinction risk, particularly in small or peripheral populations where demographic stochasticity is pronounced (Charlesworth & Willis, 2009; Frankham, 2015). Importantly, emerging evidence indicates that maintaining or restoring connectivity can facilitate genetic rescue and buffer historically bottlenecked populations against further decline (Broquet et al., 2010; Jangjoo et al., 2016). Additional to the probable effects of historical bottlenecks, this landscape has experienced extensive land-use change in the 1970s following post-partition human migration and agricultural expansion (Bahadur, 2011), further accentuating impacts of historical processes. Overall, coherent patterns with lower genetic diversity and heterozygote deficit highlight the genetic vulnerability of these populations, with low adaptive potential in the face of changing landscapes and climate, underscoring the importance of functional connectivity for their persistence.

### 4.3. Anthropogenic barriers drive genetic isolation

The extensive scatter around the almost flat IBD regression slope (Figure S4.5) suggests that geographic distance alone fails to explain the majority of spatial genetic variance within sampled individuals. Hence, we examined the effect of landscape variables in influencing gene flow with complex, non-linear landscape resistance surfaces using multi-surface optimization workflows. Landscape genetics provides a powerful framework to identify environmental resistances to dispersal (Manel et al., 2003), and to inform connectivity planning (Sharma et al., 2013).

However, connectivity research remains scarce in India’s fragmented landscapes (Thatte et al., 2020; Tyagi et al., 2024) and limited to tigers (Anwar & Borah, 2019; Biswas et al., 2022) and elephants (Neupane et al., 2022; Mukherjee et al., 2025) in the landscape.

Our study is the first to use landscape genetic approaches to understand gene flow and its drivers in the Terai. The multisurface analysis revealed agriculture cover density as the dominant barrier to gene flow in the landscape for sloth bears, explaining over 85% of the variation, followed by distance to water. Due to the strong effect of agriculture on gene flow, the high connectivity regions are primarily limited to areas with little or no agriculture (Figure 4A), scanty in the landscape. Anthropogenic features acting as significant barriers to gene flow have been revealed by multiple landscape genetic studies (Fraser et al., 2019; Yang et al., 2011). While impacts of roads and human settlements have been studied extensively across the globe (Balkenhol et al., 2020; Thatte et al., 2020; Trense et al., 2021), in the fragmented and multi-use landscapes, permeability of agricultural landscapes is a major driver of animal movement across habitat patches (Marshall et al., 2020; Niyogi et al., 2021). With human-wildlife overlap expected to increase in agricultural regions globally (Ma et al., 2024), understanding this permeability is critical for maintaining functional connectivity.

Other landscape variables, such as distance to water and vegetation indices, also influenced resistance, with areas near water and sparse vegetation showing higher resistance - consistent with studies on other bear species (Lewis, 2012; Thatte et al., 2020; Douchinsky, 2024). Counterintuitive findings for distance to settlement may reflect their skewed distribution as the sampling distribution was limited to protected areas. However, our models explained <10% of the variation in genetic distance, suggesting that other processes, such as drift or historical bottlenecks might also be important drivers of spatial genetic patterns in the landscape.

### 4.4. Genetic isolation leading to local extinction

Together these findings demonstrate the role of fragmentation with possible effects of historical processes, contributing to probable local extinction in Katarniaghat Wildlife Sanctuary (KAT) in the CTL (Yoganand et al., 2006). Unlike the contiguous metapopulations of Central India, Terai region has historically undergone rapid fragmentation due to agriculture, infrastructure, and settlement expansion (Thapa et al., 2021; Bista et al., 2024), with natural habitats now mostly confined to the bounds of protected areas. Our findings with low genetic diversity and a consistent heterozygote deficit across genetic clusters is suggestive of low population sizes and genetic erosion. Field surveys corroborate this vulnerability - KAT showed no sloth bear presence despite intensive survey effort of over 500 km. This was surprising as KAT is geographically only ∼20 km from DDW, well within species’ estimated dispersal limit of ∼13 km. This reasoning is strengthened by lack of any critical potential corridors between DDW and KAT. Although our analyses revealed transboundary connectivity across the India-Nepal border, Paudel et al. (2026) reported only two individuals in Bardia NP, Nepal adjacent to KAT, underscoring problems associated with small population sizes. With no connectivity with source populations to KAT, the latter remains isolated with possible local extinction (Yoganand et al., 2006), with limited prospects for natural demographic or genetic rescue. With most distinct genetic clustering and fragmented corridors connecting DDW with others, these populations also stand risk of population collapse in the future with increasing agriculture and loss of natural habitat.

As human modification of natural habitats intensifies, species that exhibit low tolerance to disturbance and limited permeability across agricultural and other modified matrices face the greatest risk of genetic isolation and eventual extinction, particularly in range peripheries. Apart from sloth bears, Terai hosts the northernmost populations of several large mammals, including Asian elephants, swamp deer, and fishing cats. With incongruency between genetic and geographic populations, conservation strategies must prioritize the integration of robust non-invasive genetic monitoring to identify vulnerable populations. Furthermore, targeted efforts to restore or maintain functional connectivity, ensuring that remnant populations retain their evolutionary potential and the capacity for long-term persistence is critical, especially in increasingly human dominated landscapes.

## Author Contributions

DR led project conceptualization, data collection, analyses, and manuscript writing. JCS led the data generation and co-led manuscript writing. UR assisted in conceptualization while providing critical inputs and logistical support for the project. All authors critically contributed to the drafts and approved the final version for publication. All authors were engaged in the study from the beginning to provide a broad and holistic perspective to the designing and execution of the study. The authors declare no conflicting interests.

## Data Archiving

The codes and processed VCF file can be found archived on the corresponding author DR’s <u>github</u> repository. The raw .fastq sequences can be found on the public database Indian Nucleotide Data Archive hosted by the Indian Biological Data Centre (IBDC Study Accession: INRP000770; INSDC Project Accession: PRJEB127658).

