## Supplementary Material for "Isolation to Extinction: Landscape Fragmentation and Connectivity in Sloth Bears of the Indian Terai"

This documented has supplementary data for

Associated data and codes are hosted in the primary author's GitHub repository.

#### S1. Species Identification

##### Sloth bear specific primer amplification

**Table S1: Details of primer pairs used for genetic species identification of putative sloth bear faecal DNA extracts (Thatte et al., 2018).**

| Mitochondrial Region | Primer Name | Primer Sequence | Fragment Length |
|---|---|---|---|
| Cytochrome Oxidase II | SBcoxII F | GACTGAGGACGTAGCCTC | 243 |
| | SBcoxII-1 R | TAGGAATCCCCATTATCGTG | 243 |

PCR reactions were conducted in 10ul reactions, consisting of 3ul of Master Mix, 2ul of Nuclease-free Water, 1ul of 2.5mM primer and 3ul of DNA. The reaction conditions included an initial denaturation step at 94°C for 10 minutes, followed by annealing step with 45 cycles at 94°C for 30 seconds (denaturation), 57°C for 30 seconds (annealing), and 72°C for 30 seconds (elongation), and the final extension step at 72°C for 10 min. The PCR products were visualised using gel electrophoresis to confirm for successful amplification, thereby confirming the sloth bear origin of the samples.

##### 16s primer amplification

The 16s primer (Mukherjee et al., 2016) reaction was conducted in a 10ul reaction consisting of 5ul of Master Mix, 3 ul of Nuclease Free Water, 1 ul of 2 mM primer and 1ul of DNA. The reaction conditions included an initial denaturation step at 95°C for 15 minutes (initial denaturation), followed by 45-cycle annealing step with 94°C for 30 seconds (initial denaturation), 57°C for 30 seconds (annealing), and 72°C for 30 seconds (elongation), with the final extension of 72°C for 10 minutes.

The purification prior to Sanger sequencing was done using a bead-based purification method , using magnetic beads to isolate DNA fragments of a certain size (< 300 bp) that would subsequently be sequenced. The purified products were submitted to the in-House Sequencing facility to be sequenced by Sanger Sequencing. The sequenced DNA is then matched to the online National Center for Biotechnology Information database using their BLAST feature. The sequences with over 95% match to sloth bear genetic data were positively identified as belonging to sloth bear.

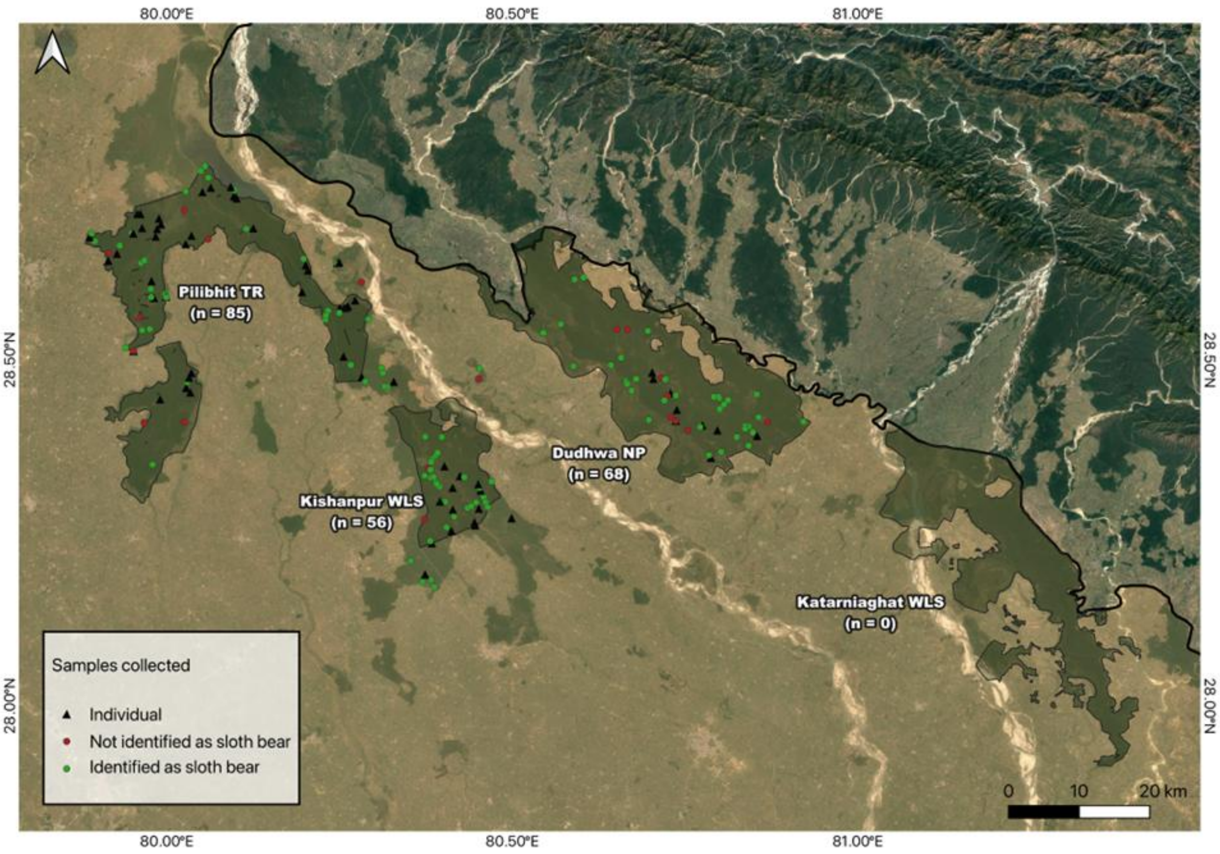

**Figure S1.1: Map of the Central Terai Landscape (Uttar Pradesh) showing sample collection sites across the four protected areas.** *The map highlights the total number of faecal samples collected at each site, the unique individuals identified, the successfully identified sloth bear samples, and those not identified as sloth bears genetically.*

### S2. DNA enrichment and library preparation

#### Faecal sample enrichment

Buffer Preparation:

1. Prepare 1X bind/wash buffer from 5X buffer stock (2.5 ml NFW + 1.2ml 5X Buffer x no. of reactions)
2. Prepare 2 M NaCl from 5M NaCl (370 µl 5 M NaCl + 630 µl 1X bind/ wash buffer)

Note: must be kept in ice.

Bead Preparation:

3. Calculate and prepare beans as required by mixing n ul of beads with n/10 of MBD-Fc protein$1 $25 ul of beads required per sample)
4. Mix beads and protein by invert mixing or mixing in a rotating mixer for 15 mins.
5. Spin down briefly and place on a magnetic stand for 2-5 minutes until the beads have collected.
6. Remove the supernatant carefully without disturbing the beads, and add 1ml of 1X wash buffer that is kept on ice.
7. Mix in a rotating mixer/ invert mix for 3 minutes.
8. Spin down and place on a magnetic stand for 2-5 minutes.
9. Repeat steps 6-8.
10. Remove the tube from the stand and add n ul of 1X wash/bind buffer and resuspend the beads by inverting or gentle pipette mixing.

Capture DNA:

11. To 32 µL of sample, add 8 µL of 5 M bind/wash buffer and 5 µL of beads.
12. Invert mix and incubate for at least 30 minutes, mixing every 5 minutes.
13. Briefly spin down and place on a magnetic stand for 2-5 minutes.
14. Remove the supernatant carefully and add 100 ul of 1X wash/bind buffer while on the stand.
15. Remove the supernatant carefully.
16. Repeat steps 13-15.
17. Wash with 1X buffer again while removing from the magnetic stand.
18. Pipette mix gently and place on the magnetic stand.
19. Discard the supernatant.
20. Add 50 ul of 2 M NaCl prepared in step 2 to each sample.
21. Pipette mix gently till suspended and incubate for 10 minutes.
22. Briefly spin down and place on a magnetic stand for 2-5 minutes.
23. Carefully transfer the supernatant to a new microcentrifuge tube.

Bead Clean-up:

24. Add 90 ul of in-house SPRI beads to the elute. (1.5X-1.8X)
25. Pipette mix and vortex for 5 minutes.
26. Spin down and place on the magnetic rack for 15 minutes.
27. Prepare 70% ethanol.
28. Discard supernatant and wash the beads twice with freshly prepared 70% ethanol.
29. Resuspend beads in 30 ul of 10 mM Tris Buffer.

Vortex, spin down and carefully transfer the elute to a fresh microcentrifuge tube.E

#### ddRAD sequencing

(Note: All calculations are for 1 reaction)

A. Sample Preparation (Extraction, Enrichment, Quantification)

1. Extract genomic DNA from fecal samples using Qiagen blood and tissue kit with Longmire's buffer.
2. Elute DNA in 200 µL AE buffer.
3. Quantify total DNA with Qubit ds-HS kit.
4. Quantify host DNA (hDNA) using qPCR with c-myc primers.
5. Concentrate samples to 20 µL with Cenrivap DNA vacuum concentrator for better hDNA capture.
6. Perform hDNA enrichment following published protocols.
7. Quantify enriched total, then concentrate enriched DNA to 10 µL for library prep.

B. Reagents Preparation (Adapters & Indexes)

8. Prepare a 10X annealing buffer (1 M Tris-HCl pH 8, 5 M NaCl, 0.5 M EDTA, water).
9. Resuspend adapter oligos to 100 µM in 10 mM Tris-HCl or TE buffer.
10. Anneal double-stranded adapters by mixing complementary oligos and heating to 97.5°C, then cooling slowly to 21°C. Store at 4°C.
11. Prepare working concentrations of adapters by diluting stocks with 10 mM Tris-HCl.
12. Create Adapter Mix combining diluted SphI and MluCI adapters with annealing buffer. Use 5 µL per reaction, store at –20°C.
13. Resuspend index primers in 10 mM Tris-HCl to 100 µM stock, dilute to 10 µM for PCR.

C. Library Construction

Restriction Digestion:

14. Prepare restriction master mix: NEB Cut Smart Buffer, SphI-HF enzyme, MluCI-HF enzyme, and water.
15. Add 10 µL of DNA to 10 µL restriction master mix (20 µL total).
16. Digest at 37°C for 3 hours, heat inactivate at 65°C for 20 minutes, then hold at 4°C.
17. Digested DNA can be stored overnight at 4°C.

Adapter Ligation:

18. Prepare ligation master mix: T4 DNA ligase buffer, T4 DNA ligase, water, and adapter mix.
19. Add 20 µL ligation mix to 20 µL digested DNA (40 µL total).
20. Incubate at 23°C for 2 hours, heat inactivate at 65°C for 10 minutes.

Clean-up of Ligation Product:

21. Add 1.5X volume of AMPureXP beads (60 µL beads to 40 µL ligation product).
22. Mix by pipetting (do not vortex), incubate 5 minutes at room temperature.
23. Place on a magnetic stand for 2 minutes and discard supernatant carefully without disturbing beads.
24. Wash beads twice with 100 µL 80% ethanol, removing ethanol completely after washing.
25. Dry beads ~1 minute (avoid overdrying).
26. Elute DNA in 20 µL 10 mM Tris-HCl pH 8.
27. Place on a magnetic stand, transfer clear elute to new tube.

Indexing PCR Amplification:

28. Prepare PCR mix: 2X NEB master mix, forward primer (i5), reverse primer (i7), and cleaned ligated DNA.
29. Total volume: 40 µL.
30. Split into four 10 µL aliquots to reduce PCR bias.
31. PCR cycling:
    1. 98°C for 30 s
    2. 16 cycles of: 98°C for 30 s, 65°C for 30 s, 72°C for 30 s
    3. 72°C for 5 min, hold at 4°C.
32. Pool four aliquots post-PCR.
33. Quantify PCR products using Qubit ds-HS kit.

Dual Size Selection Using AMPure XP Beads:

34. Add 10 µL of nuclease free water to the indexed PCR product (to 50 µL total).
35. Add 25 µL (0.5X) AMPure beads, mix well, incubate 5 min.
36. Place on magnetic stand 2 min, transfer supernatant to new tube (discard beads).
37. Add 25 µL AMPure beads to the supernatant (~100 µL total), mix and incubate for 5 min.
38. Place on a magnetic stand for 2 minutes, discard supernatant carefully.
39. Wash beads twice with 80% ethanol as above.
40. Dry beads briefly, elute final library in 20 µL 10 mM Tris-HCl pH 8.
41. Quantify final library concentration (Qubit) and check size distribution (Tapestation).

D. Library Normalization, Pooling, and Sequencing Preparation

42. Calculate library molar concentration (nM) using library size and concentration:
43. Normalize all libraries to 2 nM using 10 mM Tris-HCl pH 8.

44. Pool normalized libraries by combining equal volumes .
45. Verify pooled library concentration (~2 nM).
46. Denature and dilute pooled library according to Illumina NovaSeq 6000 guidelines.

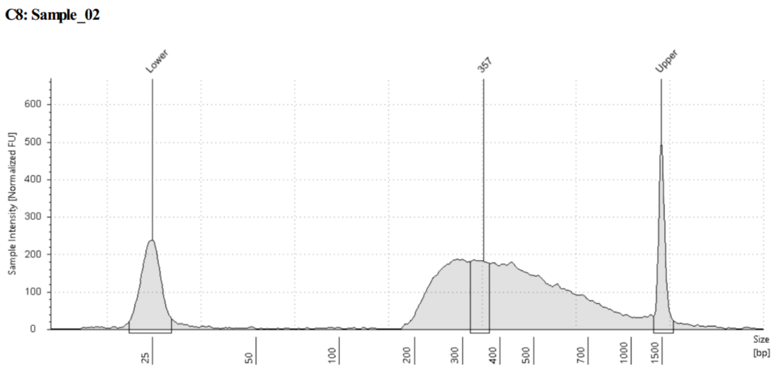

**Figure S2.1 Example ddRAD library after preparation**

### S3. Bioinformatic Analysis

**Table S3.1: Ipyrad parameter list with values and paths provided**

| PARAMETER NAME | PARAMETER DESCRIPTION | VALUES AND PATHS PROVIDED |
|---|---|---|
| assembly_name | Used to assign a prefix to all output files | sbanalysis |
| project_dir | Path to a directory where files are to be stored | slothbearanalysis |
| raw_fastq_path | Path to non-demultiplexed files | - |
| barcodes_path | Path to file containing barcodes | - |
| sorted_fastq_path | Location to demultiplexed files | /home/uramakri/utkarshschauhan/Judi/sbanalysis_filtered/inputsb/*.fastq.gz |
| assembly_method | Offers choice from either denovo clustering or reference mapping | denovo |
| reference_sequence | path to where the reference sequence is downloaded | - |
| datatype | Choice from 6 different datatypes used in RAD-seq | pairddrad |
| restriction_overhang | Detect and filter out adapter sequences | CATGC,AATT |
| max_low_quality_bases | Sets the upper limit for the number of ambiguous sites (N) allowed in reads | 5 (allows upto 5 N) |
| phred_score_offset | Reads filtered and trimmed if quality scores are not a certain value | 33 (default) |
| mindepth_statistical | Minimum depth of statistical base calling | 6 (default) |

| | | |
|---|---|---|
| mindepth_majrule | Minimum depth of majority rule base calling | 6 (default) |
| maxdepth | Sets the maximum value beyond which clusters would be excluded | 10000 (default) |
| clust_threshold | Similarity levels of two sequences being clustered together. | 0.85 (default) |
| max_barcodes_mismatch | Number of mismatches allowed | 0 |
| filter_adapters | Levels of strictness in filtering out illumina adapters | 2 (strict filtering) |
| filter_min_trim_len | Minimum length of reads after trimming to be present | 35 (default) |
| max_alleles_consensus | Highest number of unique alleles present depending on ploidy of individual | 2 (since diploid) |
| max_Ns_consens | Maximum percentage of uncalled bases allowed in a consensus | 0.05 (default ) |
| max_Hs_consens | Maximum number of heterozygous bases allowed in consensus | 0.05 (default) |
| min_samples_locus | Lower threshold for number of samples containing data at a particular locus | 4 (default) |
| max_SNPs_locus | Maximum SNPs allowed in final locus | 0.2 (default) |
| max_Indels_locus | Maximum insertions and deletions allowed in final locus | 8 (default) |
| max_shared_Hs_locus | Maximum number of shared polymorphic sites in a locus | 0.5 (default) |
| trim_reads | Specific trimming of N bases at the beginning or end of a sequence | 0, 0, 0, 0 (no extra trimming required) |
| trim_loci | Specific locus edge trimming | 0, 0, 0, 0 (no extra trimming required |
| output_formats | Choose from various output formats that can be user-defined based on type of analysis | v (can specify up to 3) |
| pop_assign_file | Assigns unique identify to samples | - |
| reference_as_filter | Used to remove sequences mapped to a reference genome | - |

### S4. Population Genetic Analysis

**Table S4.1. Comparative heterozygosity measures samples using sloth bear or sunbear reference genome for *ipyrad* based *denovo* assembly**

| **Areas Collected** | **Unique individuals** | | **$H_O$** | | **$H_E$** | | **$F_{IS}$** | |
|---|---|---|---|---|---|---|---|---|
| | **Sloth bear** | **Sun bear** | **Sloth bear** | **Sun bear** | **Sloth bear** | **Sun bear** | **Sloth bear** | **Sun bear** |

| | | | | | | | | |
|---|---|---|---|---|---|---|---|---|
| **DDW** | 7 | **10** | 0.092 | **0.083** | 0.149 | **0.137** | 0.27 | **0.27** |
| **PBT** | 43 | **41** | 0.072 | **0.075** | 0.206 | **0.206** | 0.59 | **0.58** |
| **KIS** | 16 | **15** | 0.060 | **0.061** | 0.132 | **0.123** | 0.38 | **0.36** |
| **Total** | 66 | **66** | 0.074 | **0.08** | 0.162 | **0.17** | 0.463 | **0.45** |

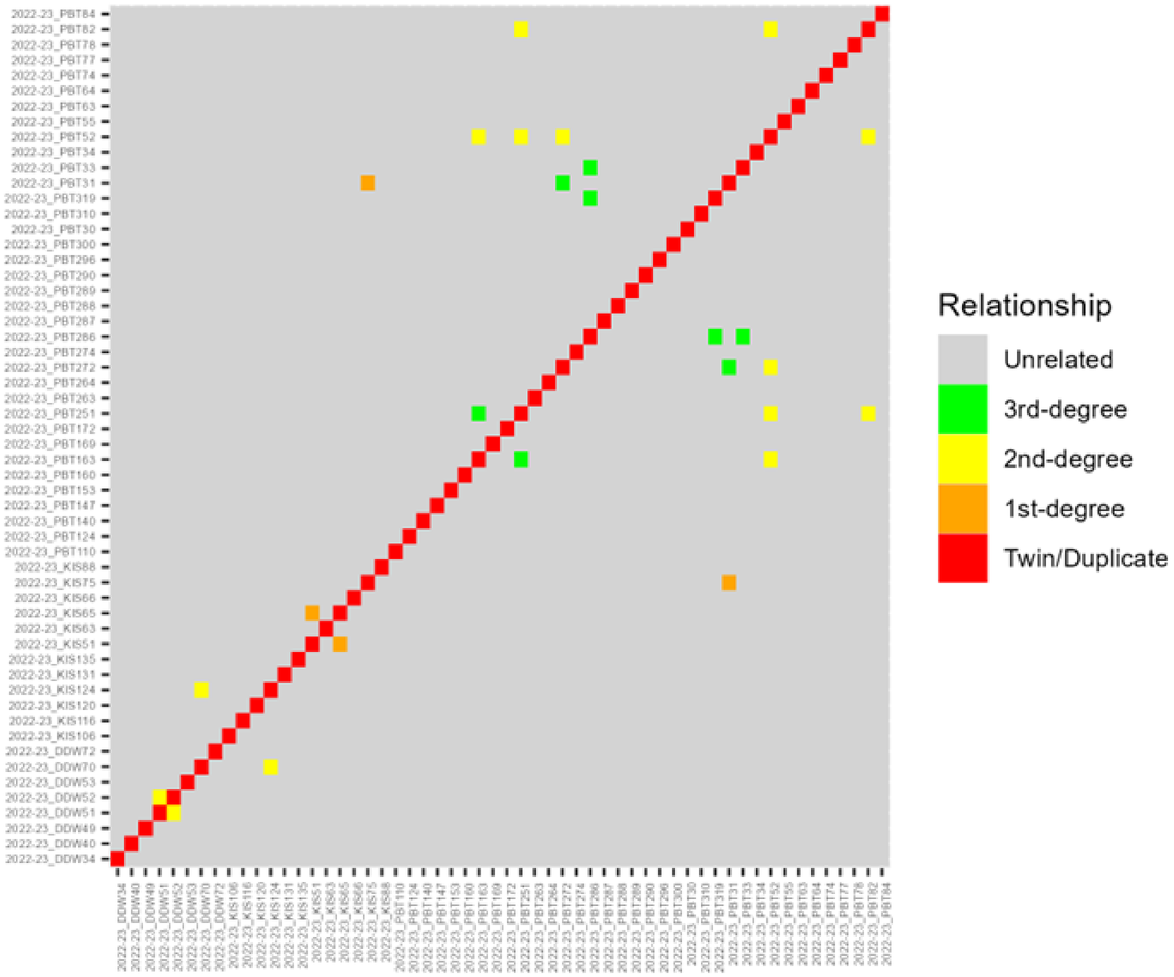

**Figure S4.1: Pairwise genetic relatedness matrix among sloth bear individuals. A pairwise relatedness heatmap plotted where each row and column represented**

**individual sloth bear samples labelled using a unique id, population (DDW, PBT, KIS) and collection year.**

**Table S4.2. Segregation of individuals with respect to their sampling location and genetic cluster with maximum membership.**

| | **K1** | **K2** | **K3** | **K4** |
|---|---|---|---|---|
| DDW | 9 | 0 | 0 | 1 |
| KIS | 0 | 2 | 2 | 11 |
| PBT | 18 | 7 | 14 | 2 |

**Table S4.3. Genetic diversity estimates across the four optimal genetic clusters.**

| **Genetic cluster** | **Ho** | **He** | **FIS** |
|---|---|---|---|
| K1 | 0.079 | 0.159 | 0.329 |
| K4 | 0.041 | 0.079 | 0.351 |
| K2 | 0.136 | 0.297 | 0.442 |
| K3 | 0.061 | 0.089 | 0.170 |

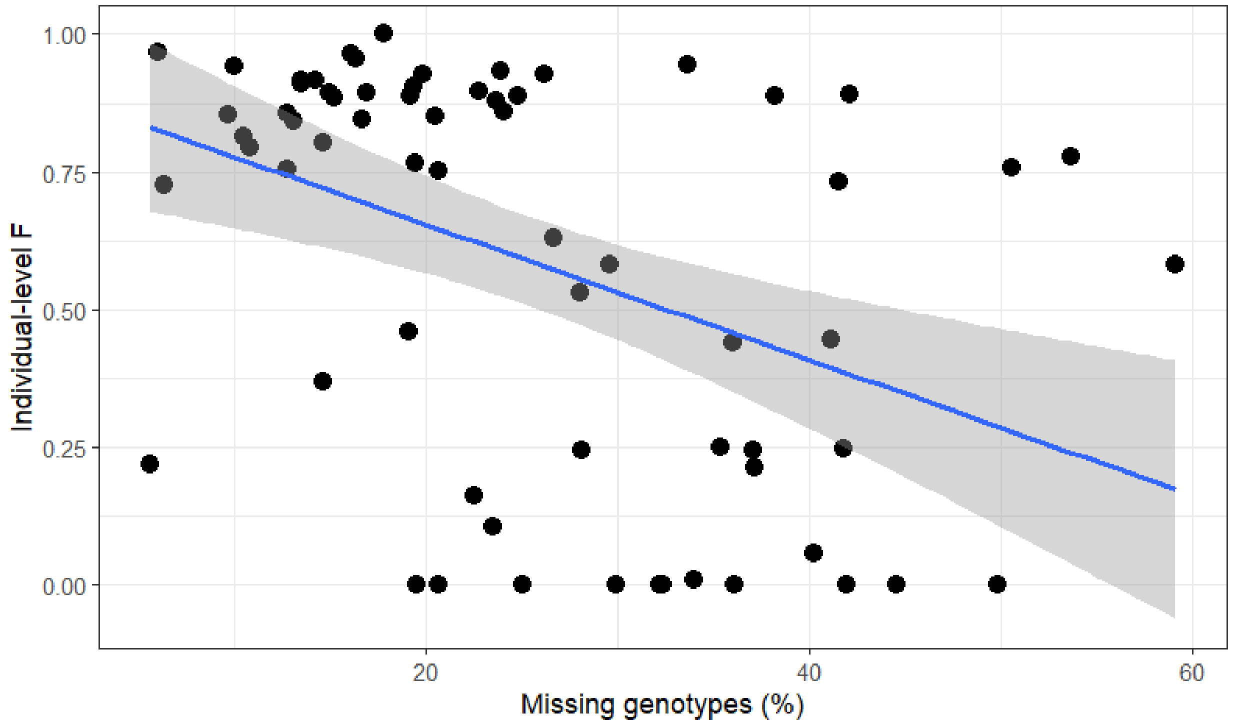

**Figure S4.3. Regression analyses investing relation between individual level inbreeding estimate and per individual genotype missingness. Spearman's ρ = −0.46, P<0.001.**

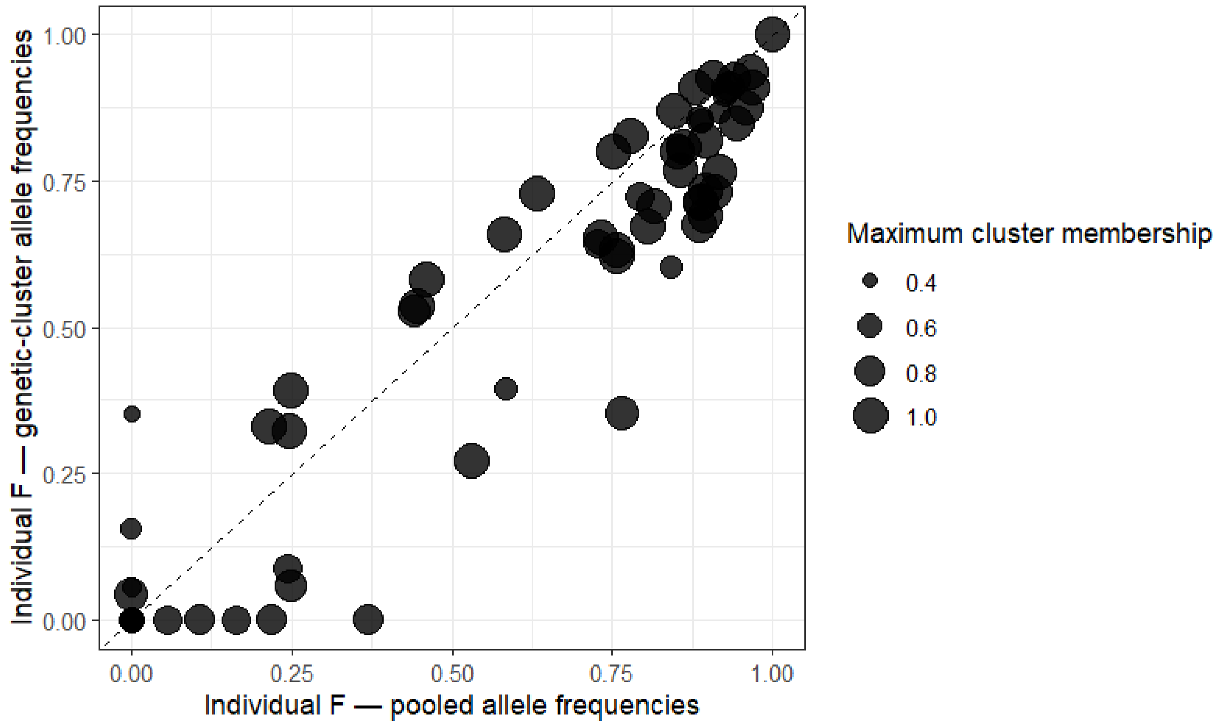

**Figure S4.4. Effect of accounting for genetic structure of individual-level F estimates with an indication of maximum cluster membership proportion for each individual. Spearman's ρ = 0.91, P<0.001.**

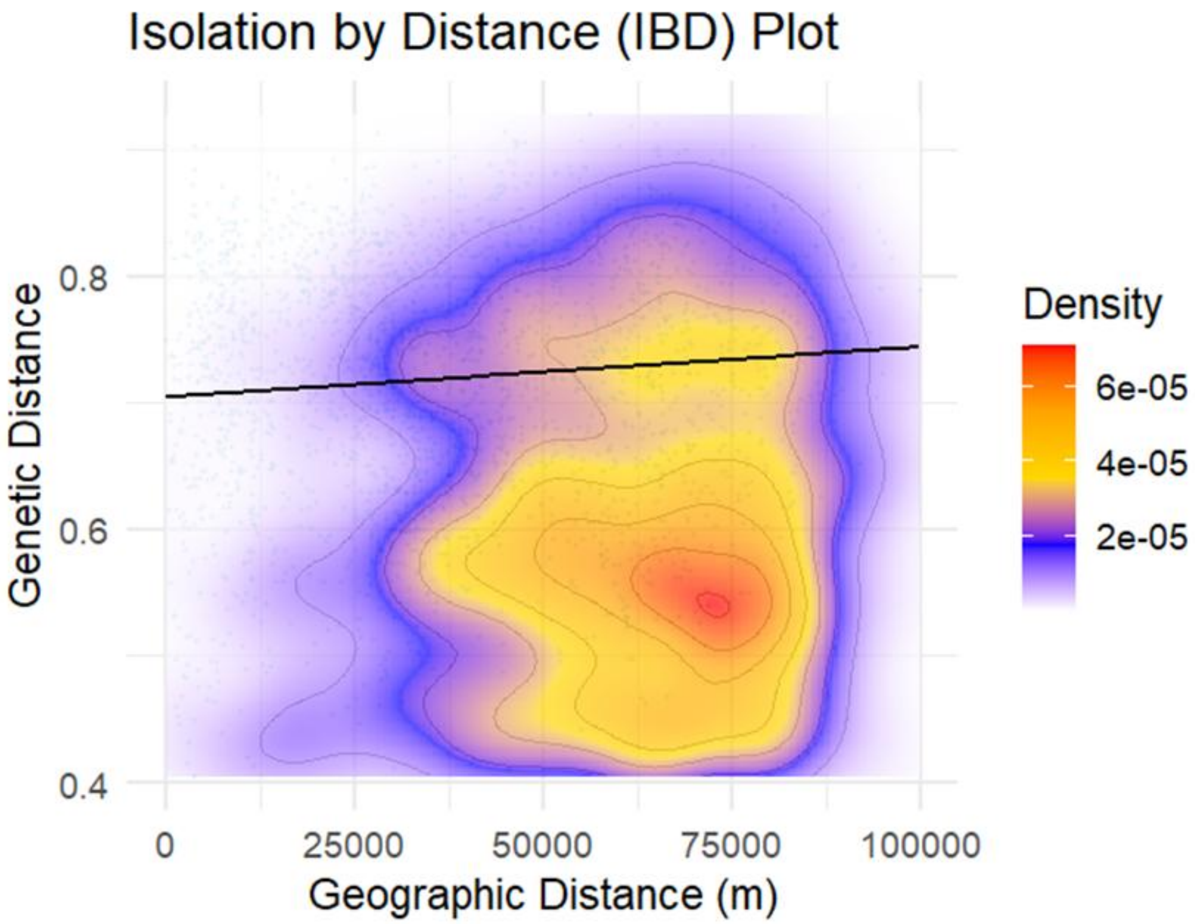

**Figure S4.5: Exploring correlation between genetic and geographic distance to detect the presence of IBD. Weak positive slope (r = 0.09; p-value < 0.042).**

### S5. Modelling functional connectivity

#### Landscape variable raster preparation

All landscape variables were generated for the study extent. To generate rasters with different scales, focal mean windows of different extents - 1 km, 2 km, 5 km, and 10 km were used using the terra package in R. All variables were processed and generated in UTM projection (EPSG:32644) to minimize transformation related errors for distance and density calculations. Uncorrelated rasters were generated for four variables - distance to water, distance to human settlement, agriculture landcover density, and enhanced vegetation index.

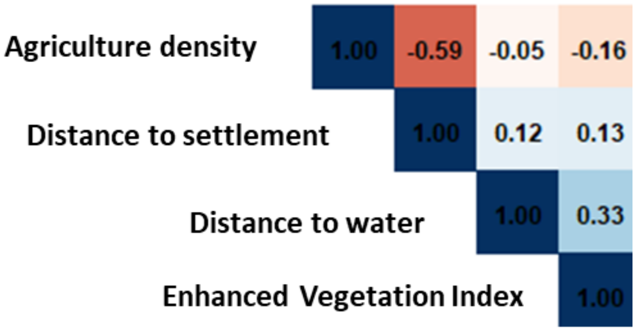

**Figure S5.1. Correlation between four landscape variables at 1km resolution**

**Distance to water**

Binary raster for inland water was generated based on category 2 of the GLCF: Landsat Global Inland Water dataset in Google Earth Engine. The binary layer was used as an input to generate the distance raster using the *"Proximity (Raster Distance)"* function in QGIS. The distance raster was resampled to match specific resolutions to create distance to water raster at different scales using the *terra* package in R.

**Distance to human settlement**

Binary raster for human settlement was generated by combining builtup categories 12-25 of the JRC Global Human Settlement Layer dataset in Google Earth Engine. The binary layer was used as an input to generate the distance raster using the *"Proximity (Raster Distance)"* function in QGIS. The distance raster was resampled to match specific resolutions to create distance to water raster at different scales using the *terra* package in R.

**Enhanced Vegetation Index**

EVI data was collated from the MODIS Terra Daily EVI dataset by averaging across year-round values for 2023. The raster scaled by a factor 0.0001 to get values ranging 0 to 1. The denser and greener the vegetation, the higher the EVI index. This rescaled raster was resampled to generate rasters for different resolutions comparable to other variable rasters.

**Agriculture landcover density**

Binary raster for agriculture land cover was generated based on category 40 of the ESA World Cover dataset in Google Earth Engine. An overlay grid of the same extent was created using the "Create Grid" tool in QGIS. The binary raster layer was used as input in the "Zonal Statistics" tool under the "Raster" menu in QGIS to calculate count and sum of the agriculture pixels for different resolutions. Then the density of agriculture land cover was calculated for each grid cell using the field calculator ((sum / count) * 100) in QGIS. Lastly, the vector density grid layer was converted to raster using the rasterize function of the terra package in R. This density raster was resampled to generate rasters for different resolutions.

#### Resistance surface optimization

Single surface optimization was performed twice for each landscape variable as recommended in the ResistanceGA manual, specifically because our optimized parameters were towards the extreme end of parameterization space. Bounds of this space were also tested twice using max.cont 1000 and 5000. Across all runs, consistent parameters were observed, and hence, in the final article values are reported for raw rasters based on a maximum parameter bound of 1000. For other shape parameters, default values as recommended in the ResistanceGA manual were used. Based on AICc values, best optimized scales for agricultural density, distance to settlement, distance to water, and EVI were 1 km, 5 km, 2 km, and 10 km respectively.

**Table S5.1: Bootstrap results of the single surface optimization**

| Surface | Scale | k | AIC | AICc | LL | Equation | shape | max |
|---|---|---|---|---|---|---|---|---|
| Agricultural density | 10 km | 4 | -8678.49 | -8678.48 | 4343.25 | Inverse-Reverse Monomolecular | 0.8520751 | 997.773 |
| **Agricultural density** | **1 km** | **4** | **-8713.11** | **-8712.45** | **4360.55** | **Inverse-Reverse Monomolecular** | **1.080398306** | **982.835** |
| Agricultural density | 2 km | 4 | -8692.61 | -8691.95 | 4350.3 | Inverse-Reverse Monomolecular | 1.122745553 | 890.089 |
| Agricultural density | 5 km | 4 | -8682.3 | -8681.64 | 4345.15 | Inverse-Reverse Monomolecular | 1.005633142 | 979.142 |
| Distance to settlement | 10 km | 4 | -8650.35 | -8650.35 | 4329.18 | Monomolecular | 0.9831576 | 990.215 |
| Distance to settlement | 1 km | 4 | -8647.5 | -8646.85 | 4327.75 | Distance | 9 | 9 |
| Distance to settlement | 2 km | 4 | -8657.59 | -8656.93 | 4332.79 | Monomolecular | 0.70980208 | 875.393 |
| **Distance to settlement** | **5 km** | **4** | **-8658.59** | **-8657.93** | **4333.29** | **Monomolecular** | **0.753467284** | **955.767** |
| Distance to water | 10 km | 4 | -8664.72 | -8664.71 | 4336.36 | Inverse Monomolecular | 1.587101 | 995.707 |
| Distance to water | 1 km | 4 | -8670.5 | -8669.85 | 4339.25 | Inverse Monomolecular | 1.51350382 | 980.641 |
| **Distance to water** | **2 km** | **4** | **-8673.35** | **-8672.69** | **4340.67** | **Inverse Monomolecular** | **1.492445851** | **796.534** |
| Distance to water | 5 km | 4 | -8673.31 | -8672.66 | 4340.66 | Inverse Monomolecular | 1.515183482 | 972.644 |
| **Enhanced Vegetation Index** | **10 km** | **4** | **-8665.09** | **-8664.43** | **4336.54** | **Inverse Monomolecular** | **5.12087645** | **999.965** |
| Enhanced Vegetation Index | 1 km | 4 | -8654.92 | -8654.27 | 4331.46 | Inverse-Reverse Monomolecular | 2.355771073 | 975.683 |
| Enhanced Vegetation Index | 2 km | 4 | -8653.53 | -8652.87 | 4330.76 | Inverse-Reverse Monomolecular | 3.319006792 | 998.147 |
| Enhanced Vegetation Index | 5 km | 4 | -8649.72 | -8649.07 | 4328.86 | Monomolecular | 6.339855403 | 998.769 |

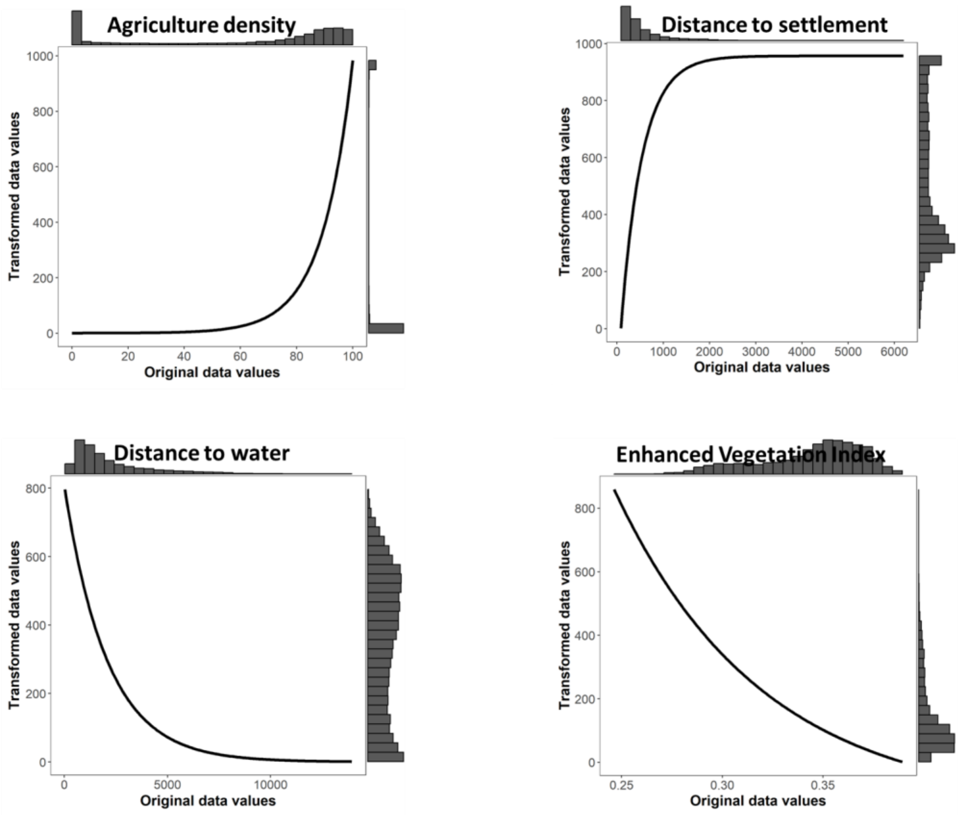

**Figure S5.2: Single surface resistance optimization for rasters at respective optimized scales**

**Table S5.2: Multisurface optimization results across all variable combinations. Agriculture density is the most important variable in all the top model combinations.**

| subset | model | k | AIC | AICc | R2m | R2c | logLik |
|---|---|---|---|---|---|---|---|
| 2 | agri_dens_1km.asc+dist_water_30m_utm_2km.asc | 3 | -8709.5 | -8709.12 | 0.100556 | 0.581532 | 4357.752 |
| 1 | agri_dens_1km.asc+dist_settlement_10m_utm_5km.asc | 3 | -8709.49 | -8709.11 | 0.100912 | 0.581938 | 4357.746 |
| 3 | agri_dens_1km.asc+MODIS_EVI_rescaled_500m_utm_10km.asc | 3 | -8709.41 | -8709.03 | 0.100449 | 0.581496 | 4357.707 |
| 8 | agri_dens_1km.asc+dist_settlement_10m_utm_5km.asc+MODIS_EVI_rescaled_500m_utm_10km.asc | 4 | -8701.09 | -8700.43 | 0.086543 | 0.570472 | 4354.545 |
| 7 | agri_dens_1km.asc+dist_settlement_10m_utm_5km.asc+dist_water_30m_utm_2km.asc | 4 | -8701.06 | -8700.41 | 0.086697 | 0.570681 | 4354.532 |
| 9 | agri_dens_1km.asc+dist_water_30m_utm_2km.asc+MODIS_EVI_rescaled_500m_utm_10km.asc | 4 | -8701.05 | -8700.39 | 0.086304 | 0.57021 | 4354.524 |
| 4 | dist_settlement_10m_utm_5km.asc+dist_water_30m_utm_2km.asc | 3 | -8675.27 | -8674.88 | 0.137065 | 0.561871 | 4340.634 |
| 6 | dist_water_30m_utm_2km.asc+MODIS_EVI_rescaled_500m_utm_10km.asc | 3 | -8675.27 | -8674.88 | 0.138706 | 0.562801 | 4340.634 |
| 10 | dist_settlement_10m_utm_5km.asc+dist_water_30m_utm_2km.asc+MODIS_EVI_rescaled_500m_utm_10km.asc | 4 | -8673.28 | -8672.62 | 0.145728 | 0.569353 | 4340.639 |
| 11 | agri_dens_1km.asc+dist_settlement_10m_utm_5km.asc+dist_water_30m_utm_2km.asc+MODIS_EVI_rescaled_500m_utm_10km.asc | 5 | -8671.11 | -8670.11 | 0.053843 | 0.541719 | 4340.555 |
| 5 | dist_settlement_10m_utm_5km.asc+MODIS_EVI_rescaled_500m_utm_10km.asc | 3 | -8667 | -8666.61 | 0.110133 | 0.549954 | 4336.5 |

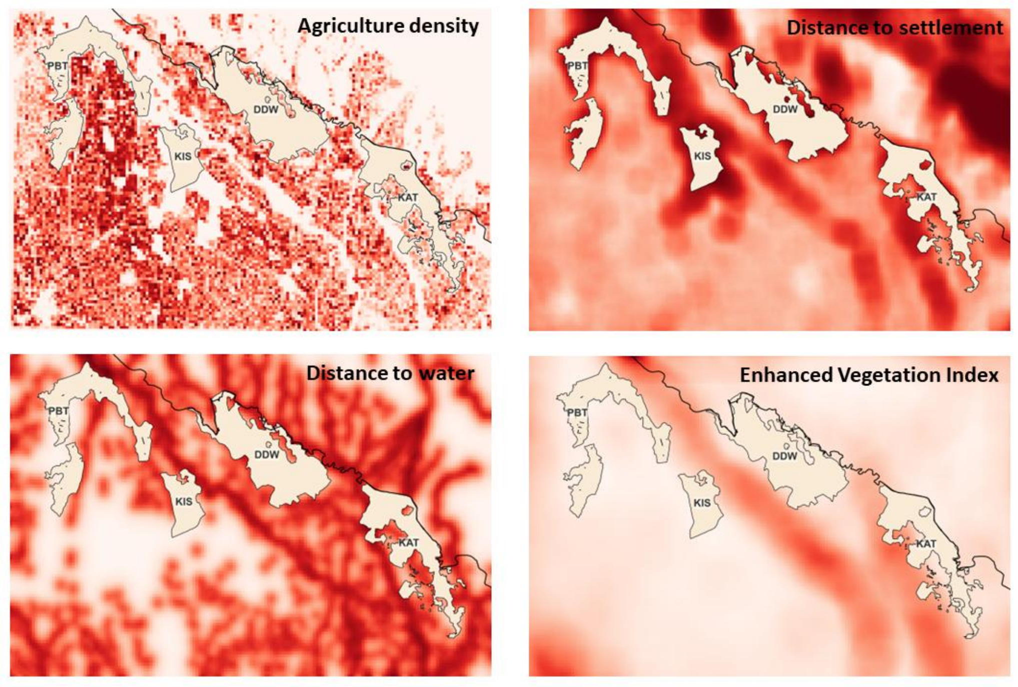

**Figure S5.3: Single variable scale optimized resistance surface across study region (raster resolution = 1 km). Darker colours represent higher resistance.**

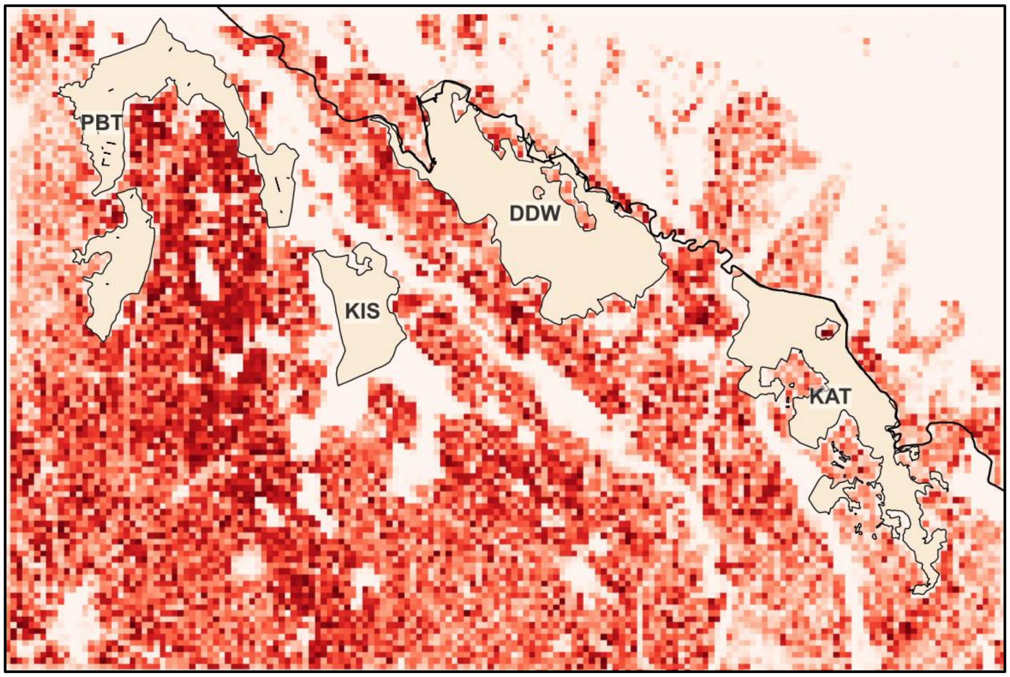

**Figure S5.4: Composite multi-scale resistance surface across study region. Darker colours represent higher resistance. This composite surface consists of contributions from agricultural density (86.11%) and distance to water (13.89%).**
